# Locomotor adaptation can persistently reorganize stride-to-stride regulation of centre-of-mass error dynamics

**DOI:** 10.64898/2026.07.10.737847

**Authors:** Daphna Raz, Rachel Marbaker, Sriram Sankaranarayanan, Alaa A. Ahmed

## Abstract

Locomotor learning in novel environments relies on a gradual alteration of motor output to achieve a desired state. Changes in gait outcomes such as step-length asymmetry are typically used to describe this process. Recently, stability-relevant adaptations, quantified by scalar metrics such as the margin of stability, have also attracted interest. Yet humans, in part, regulate walking stability from stride-to-stride. Scalar metrics only provide instantaneous snapshots of stability and fail to capture rules governing fluctuations from one stride to the next. Here, we investigate whether locomotor adaptation changes how movement regulation evolves *across* strides using a multidimensional, dynamical systems model of centre-of-mass (CoM) based locomotion error. We apply this approach to split-belt locomotor adaptation, where participants walk on a treadmill with a separate belt for each foot. One belt moves faster than the other, driving participants to adapt compensatory gait patterns due to asymmetry. Using our stride-to-stride model, we find that, in addition to reducing CoM error while adapting, multidimensional error regulation dynamics are also adapted. This structural adaptation persists upon re-exposure to split-belt conditions. Our findings show that split-belt locomotor adaptation includes an adaptation of the structure of stride-to-stride CoM movement regulation and that this structure may be rapidly recalled.

## 1 Introduction

Despite the inherent instability of bipedal locomotion, humans are able to adapt to walking in a variety of novel environments. They traverse sandy beaches, hike on rocky trails, and navigate icy winter sidewalks, often changing their gait mechanics without falling. Understanding the process by which humans manage such feats is not only of scientific interest, but could contribute to the design of effective balance rehabilitation protocols and improve our understanding of the heightened fall risk that occurs with certain movement disorders, such as multiple sclerosis or Parkinson’s. Although situations requiring gait adaptation are ubiquitous in daily life, systematically studying real-world examples of locomotor adaptation is difficult due to factors such as limited sensing capabilities and high environmental variability. A complete understanding of how humans learn to locomote while maintaining stability thus remains out of reach.

Split-belt treadmill walking has emerged as a popular, lab-based approach to study locomotor adaptation. Participants walk on a two-belt treadmill, with each leg on a separate belt. Under nominal conditions, referred to as ‘tied-belt’, the belts move at the same speed. During ‘split-belt’, one belt of the treadmill moves up to 4 times faster than the other belt, creating inherently asymmetrical and unnatural dynamics [1], [2], [3], [4], [5]. The majority of studies focus on spatiotemporal kinematics and how they change upon exposure to the split belt perturbation. A typical response to this novel motor-adaptation task is that the magnitude of step-length asymmetry rises consistently and sharply upon first exposure, gradually decaying close to baseline over time [6], [7], [8]. Because most participants return to a fairly symmetric baseline, i.e. close to zero step-length asymmetry, this signal is interpreted as an error that is progressively corrected during adaptation [8], [9]. Upon re-exposure, the decay in asymmetry occurs at a faster timescale, providing evidence of recall [2], [7], [10], [11], [12].

While outcomes such as reduction in step-length asymmetry provide valuable insight into high-level processes governing locomotor learning, they cannot explain how humans adapt while simultaneously remaining upright. Stability adaptations during split-belt walking have only recently attracted attention. Several studies have quantified stability-related changes using scalar metrics. These stability metrics include whole body angular momentum, the sum of body-segment angular momenta about the body CoM, [13], or more commonly, the mediolateral Margin of Stability (MoS) [14], [15], [16], [17], a stability margin consisting of a linear combination of position and velocity of the CoM relative to the base of support [18]. Mediolateral MoS rises and then converges during split-belt adaptation in both legs [15], with convergence occurring at a faster time scale than reductions in step-length asymmetry [14]. Recent studies further suggest that suboptimal adapted movement patterns are in part explained by minimizing fall risk [19], and that stability drives early adaptation [14], [20].

These results provide strong evidence that stability-relevant adaptations occur during split-belt adaptation but leave many open questions. It is unclear whether adaptation changes only observable stability metrics, or whether it also changes latent structure governing how stability is regulated. Analysing decay rates implicitly treats MoS as an error signal analogous to step-length asymmetry, suggesting that humans learn to reduce stabilization error to some desired amount. While MoS does appear to converge toward a more stable steady state, this raises the question of which reference state is being targeted, and whether error reduction alone is a sufficient explanation. Rather, stability error reduction may be a manifestation of the adaptation of a more complex latent control process. Humans regulate walking throughout the gait cycle, notably from stride to stride [21], [22]. An alternative possibility is therefore that the nervous system not only reduces stability error but also changes the dynamics of how this error is regulated across strides. Metrics such as MoS only provide an instantaneous snapshot of stability and cannot resolve this question.

As a consequence of our limited understanding of stability and adaptation, we also do not know whether stability-related adaptations can be retained and recalled. Most stability-focused studies do not investigate savings, where participants are reexposed to the split-belt perturbation. This limits the conclusions that can be drawn about what precisely is being learned with respect to stability, offering insight only into the adaptation process itself, not what is possibly recalled from that process. Recent work has suggested that split-belt adaptation may involve the rapid recall of previously learned movement mappings beyond scalar error reduction [23]. Investigating the evolution of stability upon re-exposure to the perturbation, would enable the teasing out of whether stability-related ‘error’ is simply reduced faster upon re-exposure, or whether there is a persistent, recalled change in how the dynamics of the CoM are regulated.

To address these open questions, we leverage approaches that are grounded in dynamical systems models of walking. Prior work has demonstrated that CoM state and subsequent foot placement are tightly correlated across the mediolateral and sagittal plane [24], and that simple models of walking exhibiting a human-like reactive perturbation response can be fit from walking data [25]. We build on these methods to develop a model of stride-to-stride CoM error. Our model simultaneously accounts for CoM state in three dimensions, rather than representing a single outcome in one plane over time, as is typical with scalar stability metrics. We use walking data to fit a linear model of stride-to-stride CoM dynamics that defines how CoM error state at the current stride predicts the state at the next stride. This gives us a dynamical structure that represents CoM regulation not just at an instant in time, but rather as system, and provides us with analytical tools from linear systems theory.

We apply this simple walking model [25] to a dataset of 14 participants undergoing a split-belt treadmill experiment including baseline, learning (adaptation), washout (deadaptation), and savings (readaptation) blocks. Formulating a CoM error signal relative to the states of the model, we examine both how this error signal is reduced within blocks and how the stride-to-stride dynamics governing its reduction differ across blocks. This leads us to the following hypotheses:

**(H1)** Humans reduce CoM error during split-belt adaptation

**(H2)** Split-belt adaptation reorganizes the structure of stride-to-stride CoM movement regulation

**(H3)** Early savings reflects recall of previously learned structure.

We first address H1, comparing early and late CoM error within experimental blocks. Establishing that CoM error decays in a manner suggestive of an adaptive, learned process provides the necessary foundation for H2, a stride-to-stride analysis characterizing the structure of this error regulation.

We test H2 by learning a linear system from our dataset connecting the CoM state at one stride to the state at the next stride. This linear system approximates how the nervous system corrects errors from stride-to-stride. The systems are learned from steady-state data and reflect the baseline error dynamics unique to each block and each leg. We analyse the temporal and geometric structure of these systems, using the eigenvalues and eigenbases to compare how CoM error dynamics differ across walking conditions.

Finally, we investigate whether any learned structure is reflected in the early strides of savings (H3). We evaluate how well stride-to-stride CoM error correction during early savings can be explained by the steady-state dynamics established during the learning block, accounting for the shared dynamics of the split-belt treadmill. During early savings, participants are re-adapting after washout. If early savings is aligned with steady-state learning dynamics, this implies that error reduction during early savings is not simply an accelerated version of the learning process, nor are the error dynamics driven purely by the mechanics of the split-belt treadmill. Rather, participants are rapidly recalling a previously learned structure.

## 2 Methods

We first define a CoM error state based on a simple dynamic walker, deriving a three-dimensional linear model of stride-to-stride error dynamics (Section 2.1). Using a previously collected split-belt treadmill dataset (Section 2.2), we compute stride-to-stride CoM error trajectories, test for CoM error reduction, and fit block-specific linear models (Sections 2.3–2.4). We then compare the resulting dynamics using their eigenstructure (Section 2.5) and investigate whether learned dynamics are recalled during savings (Section 2.6).

### 2.1 Walking Model and CoM Error

We model the human as a simple 3D inverted pendulum with mass concentrated at a single point. The full state of the CoM consists of position and velocity in the mediolateral (*x*), anteroposterior (*y*), and vertical directions (*z*). A step is initiated by an axial impulse at pushoff of the trailing leg followed by heel strike on the leading leg (Figure 1).

**Figure 1:**
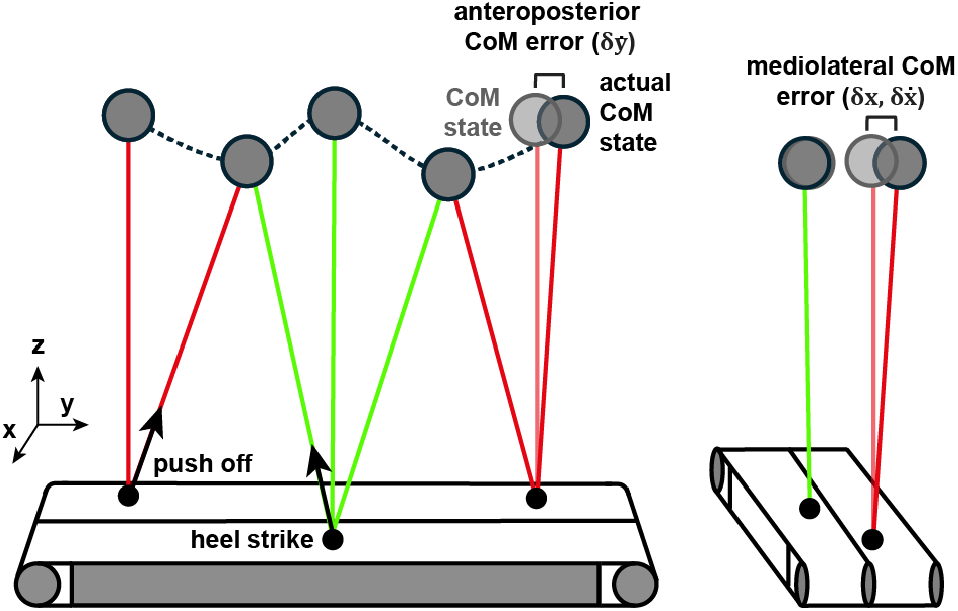
Simple inverted pendulum model of split-belt walking, with CoM error represented in the sagittal (left) and frontal plane (right). Faded pendula represent the nominal CoM state at midstance and solid colours represent the actual state. The CoM error signal is the difference between the nominal and actual CoM states. Figure adapted from [25].

Rather than model within-stride dynamics explicitly, we analyse closed loop stride-to-stride dynamics on a Poincaré section defined at midstance, or the instant when the CoM is directly above the stance foot in the sagittal plane. This reduces periodic nonlinear dynamics to a discrete linear map that records the state each time it crosses a predefined surface [26]. Assuming a fixed leg length, midstance CoM height is constant, eliminating vertical position and velocity as free states. Furthermore, midstance anteroposterior position is fixed by definition, leaving three free states: mediolateral position *x*, mediolateral velocity 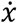, and anteroposterior velocity 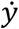.

Given these reduced dynamics, we define a reference state at midstance as the nominal CoM state, approximately corresponding to the CoM being centred over the stance foot with negligible mediolateral and sagittal-plane velocity relative to the belt speed (Figure 1). Deviations from this reference evolve according to the stride-to-stride map

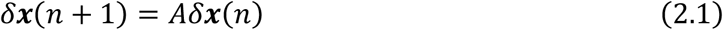

where *δ****x***(*n*) is the vector of CoM error states 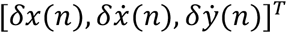 and *n* indexes the successive crossings of the Poincaré section per stride (see [25] for a detailed derivation).

### 2.2 Experiment and data processing

We apply our framework to the control group of a previously published split-belt treadmill experiment [27]. Pelvic (left/right anterior and posterior superior iliac spines) and ankle (left/right malleolus) marker trajectories from 14 participants (9 female, age 24 ± 3.6 years, weight 71 ± 15.2 kg, height 1.64 ± 0.13 m) were analyzed. Participants wore an unweighted vest as part of the original study design.

After a short acclimatization, participants completed six walking blocks (Figure 2), with short pauses between blocks: fast baseline, slow baseline, learning, washout, savings, and final washout. We use data from the first through fifth block, starting with the fast (1.5 m/s belt speed) and slow (0.5 m/s belt speed) baseline blocks, lasting two minutes each. The learning block followed, where the split-belt perturbation (belt ratio 3:1 with speeds at 1.5 m/s and 0.5 m/s) was introduced and remained constant for 10 minutes while participants walked continuously. Learning was followed by washout, a 15-minute block of continuous walking with tied-belts at the slow speed. During savings, the belt speeds split again for 10 minutes, keeping fast leg consistent with the first learning block (experiment description adapted from [27]).

**Figure 2:**
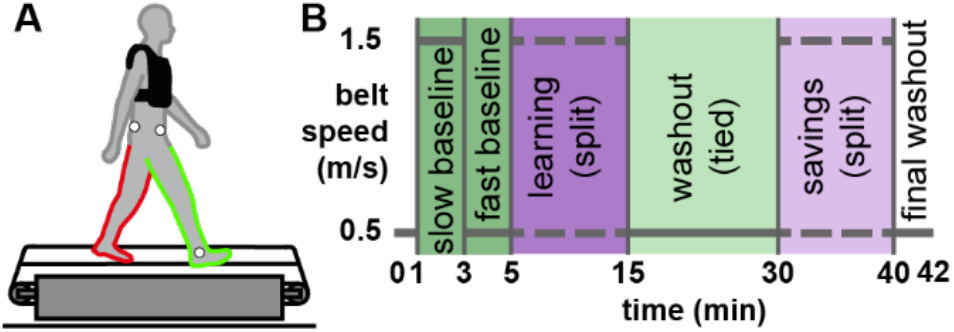
Adapted figure from [27] describing the split-belt experiment. A: Experimental Setup. We used a limited lower body marker set consisting of two pelvic and one ankle marker per side. B: Experimental Protocol. The protocol consisted of alternating tied-belt (green) and split-belt (purple) blocks.

Our modelling framework is structured around the dynamics of CoM errors at midstance 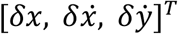 which are not directly measured in the experiment. We approximate the CoM position as the average of the four pelvic markers (left/right anterior and posterior superior iliac spines), with CoM velocity obtained by numerical differentiation. The ankle marker position and velocity define the reference state at midstance, and CoM error is computed as the difference between the ankle and CoM position and velocity (Supplement S2).

### 2.3 Investigating scalar CoM error reduction

Although the overarching goal is to better understand how the nervous system corrects errors from stride-to-stride, we must first establish that CoM error is reduced during split-belt adaptation (H1). If CoM error (*δx*) is relevant for adaptation, we expect its magnitude would increase with perturbation, i.e. with initiation of split-belt walking, and decrease with learning. We operationalize this error as the CoM midstance error defined based on the model states from the previous section: 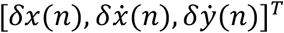. We test H1 using repeated measures ANOVA and planned comparisons as follows.

We compute the average error of the first four and last ten strides from each block. For the slow leg, we compare early and late error in baseline, learning, washout, and savings. We make the same comparisons for the fast leg, with the exception of washout, where there is no fast belt. To determine whether CoM error changes significantly during the course of the experiment, we conduct a repeated measures ANOVA with all early and late block samples as levels, with Holm-Bonferroni correction for multiple comparisons (*α* = 0.05). We perform the RM-ANOVA separately for each leg and each state variable. We follow the RM-ANOVA with the planned comparisons between early and late epochs within each block, per leg and per state.

### 2.4 Computation and validation of CoM error model

After analysing the behaviour of the CoM error per dimension from a temporal perspective, we fit the three-dimensional linear map defined in Equation (2.1) from the same dataset, discarding the initial transient phase and using steady-state data only. For each block, we compute the matrix *A*_block_, where block ∈ {B,L,W,S}. Subscripts B, L, W, and S refer to the experimental block labels Baseline, Learning, Washout, and Savings and will be used throughout the text. Solving for *A*_block_ amounts to a least squares problem (Supplement S3).

After computing *A*_block_, we confirm that the resulting model is plausible by checking that the linear dynamics are stable and that it has reasonable predictive power by computing the normalized root mean squared error of the model prediction (details and results in Supplement S6).

We also establish that the model captures meaningful multi-stride dynamical structure by computing a skill score [28]. The skill score is a method for comparing the mean squared error (MSE) of a model with a desired benchmark model, and can be interpreted as a ‘percent improvement.’ A positive skill score means that the model performs better than the benchmark comparator, while a negative skill score indicates worse performance. We compare our model to a persistence model, where the predicted next state is simply the current state, and a mean predictor model that always predicts the steady-state mean. We compute skill scores relative to both models over multiple steps, for each leg, block, and state separately:

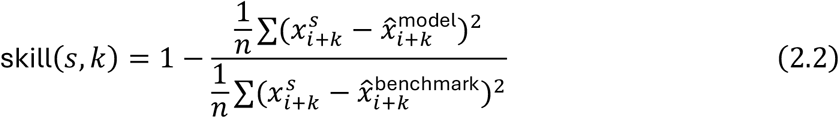

where *s* is the state, *k* is the number of strides and *n* = *N*_samples_ − *k* is the number of valid stride pairs at horizon *k*.

One-step skill relative to the mean predictor is equivalent to *R*^2^. For a stable system, skill relative to the mean predictor model should eventually decay to zero. This is not necessarily true for skill relative to the persistence model. Here, we expect a general decline in performance as our model is identified using single-stride transitions and is not meant to capture the long-term autocorrelations that are present in human walking. Thus, with increasing *k*, we expect both benchmark models to eventually outperform our model, however we prioritize performance at the first stride (*k* = 1).

We include the mean predictor model as this is a familiar benchmark. However, the persistence model is a more meaningful and difficult benchmark for noisy time series data [29]. This is particularly true in the case of nominal human gait, which exhibits long-range statistical persistence [21], implying that the current state is already fairly predictive of the next state. By construction, our Poincaré section model does not explicitly capture higher order temporal dependencies. Thus, any improvement over the persistence model indicates dynamical structure at the stride-to-stride level beyond local autocorrelation. We interpret a skill score relative to the persistence benchmark of greater than 10% as sufficient evidence of meaningful predictive structure, with performance over longer time horizons indicating the models’ ability to capture slower evolving dynamics as well.

### 2.5 Comparing steady-state dynamics between blocks

We next test whether split-belt adaptation reorganizes the steady-state dynamics of CoM regulation (H2). As linear systems are well-described by their dominant modes, our approach is to compare the eigenstructure (eigenvalues and eigenvectors) between blocks.

Due to the small sample size, we test block-level hypotheses via bootstrapping. We fit *A*_block_ over 2000 bootstrap iterations, sampling by participant with replacement, and computing the 95% confidence intervals empirically as the 2.5^th^ and 97.5^th^ percentiles. At each iteration, we compute the eigenvalues for each *A*_block_, using the resulting bootstrapped confidence interval to confirm that the estimated dynamics are consistently stable and to identify differences in stability between legs and blocks. This establishes that there is no loss of stability across the bootstrapped distribution, determines whether stability is lower in split-belt blocks, and supports our main analysis using eigenvector direction as the primary indicator of changes in stride-to-stride steady-state error dynamics.

To determine whether split-belt adaptation reorganizes the steady-state dynamics of CoM regulation (H2), we test whether the dynamics of the learning and savings blocks are more similar to each other than to the baseline block. We represent the dynamics via the dominant eigenvector. For matrices *A*_1_and *A*_2_ with dominant eigenvectors ***e***_1_ and ***e***_2_, we define similarity as

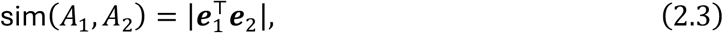

which is a cosine similarity clamped to [0,1]. Our contrast statistic is

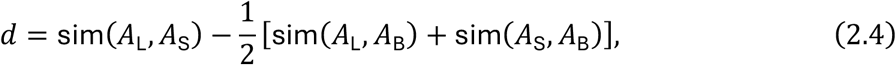

where subscripts L, S, and B denote learning, savings, and baseline respectively. A positive *d* indicates that learning and savings are more similar to each other than either is to baseline. We reject the null hypothesis at *α* = 0.05 if the bootstrapped 95% confidence interval of *d* is positive and excludes zero.

For the slow leg only, we additionally test whether split-belt and tied-belt conditions form distinct clusters via a within-between contrast. Defining within-condition similarities as sim(*A*_L_, *A*_S_) and sim(*A*_B_, *A*_W_) for split-belt and tied-belt respectively, and between-condition similarities as all pairwise split-belt vs. tied-belt comparisons, our contrast statistic is

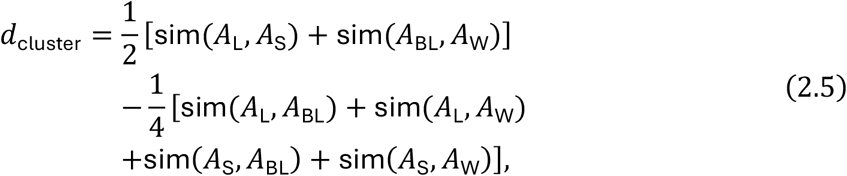

where a positive *d*_cluster_ indicates stronger within-condition than between-condition similarity.

### 2.6 Analysing recall during savings

To explore whether there is any evidence of retention of learned dynamics in early savings (H3), we compare the predictive capabilities of the learning dynamics, which reflect the same experimental conditions as savings, to the washout dynamics, which are the most recently experienced dynamics prior to the savings block. We are interested in understanding whether any of the underlying dynamical structure is retained.

We test which model (learning or washout) better predicts early savings by computing single-step skill scores (Equation 2.2) over the first 30 strides of the savings data, across participants. Rather than comparing to benchmark models, we compare the CoM error models for each block against each other:

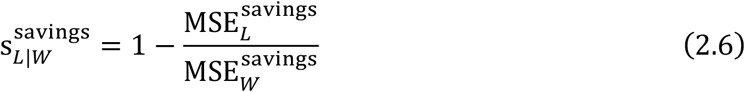

As *A*_L_ and *A*_W_ are computed relative to the block mean, we express the early savings data in the correct block coordinates by subtracting the mean of each block. If the early savings error is better explained by *A*_L_ than *A*_W_, then the skill score 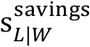 should be positive.

Because savings error dynamics may be immediately imposed by the constraints of the split-belt treadmill, we compare how well *A*_L_ predicts early learning with how well it predicts early savings:

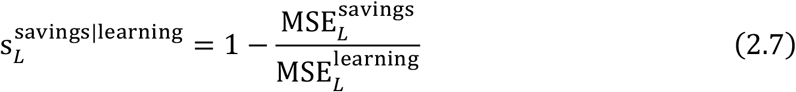

While this is not a skill score in the formal sense and cannot definitively indicate whether dynamics are truly learned, we are interested in the relative improvement. If 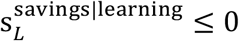, *A*_L_ would seem to predict early savings and early learning equally well or predict early learning better.

On the other hand, a positive skill score 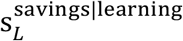 would indicate that *A*_L_ better predicts early savings than early learning. This would provide evidence that the predictive ability of *A*_L_ relative to *A*_W_ for early savings error is not explained solely by the split-belt treadmill itself. Accordingly, this would suggest that learned dynamics develop over the course of the learning block, to be quickly recalled within the early strides of savings.

To ensure that the fit is not dominated by outliers, we compute these statistics over 2000 bootstrap iterations, resampling by participant. We emphasize that these comparisons are out-of-sample by design, as *A*_L_ and *A*_W_ are estimated exclusively from steady-state data.

## 3 Results

We establish that participants significantly reduce CoM error during learning. This indicates that CoM error is a meaningful learning signal that rises with perturbation and later decays, supporting H1. After transient error reduction, the steady-state linear error dynamics differ significantly between split-belt and tied-belt blocks, supporting H2. The steady-state dynamics of the learning block predict error trajectories during the first 30 strides of savings significantly better than the washout dynamics, and also predict early savings significantly better than early learning, supporting H3 and indicating the possibility of structural recall.

### 3.1 Scalar CoM error is reduced during learning and washout

CoM error exhibited significant changes over the course of the experimental blocks. for each leg and each state (RM-ANOVA, all p values < 1 × 10^−3^). In the learning block, error was reduced from early learning to late learning in all three states and both legs (all p values < 4 × 10^−3^), save for anteroposterior velocity for the slow leg ( p = 0.343). In the washout block, where error was computed for the slow leg only, all states showed reductions from early washout to late washout (all p values < 1 × 10^−3^). In savings, where learning is most rapid, mediolateral position error was reduced for both legs, with mediolateral velocity reducing only for the slow leg, and no reductions detected for anteroposterior velocity (full test statistics in Supplement S7).

In the learning block, the raw data plots (Figure 3) show a consistent initial overshoot in the lateral (positive) direction for *δx* in both legs. The mediolateral velocity error is initially positive for the slow leg and negative for the fast leg, indicating that the CoM motion is directed toward the slow leg. The large initial (negative) anteroposterior velocity error in both legs eventually converges, meaning that the CoM progresses forward faster relative to the stance foot than during steady-state.

**Figure 3:**
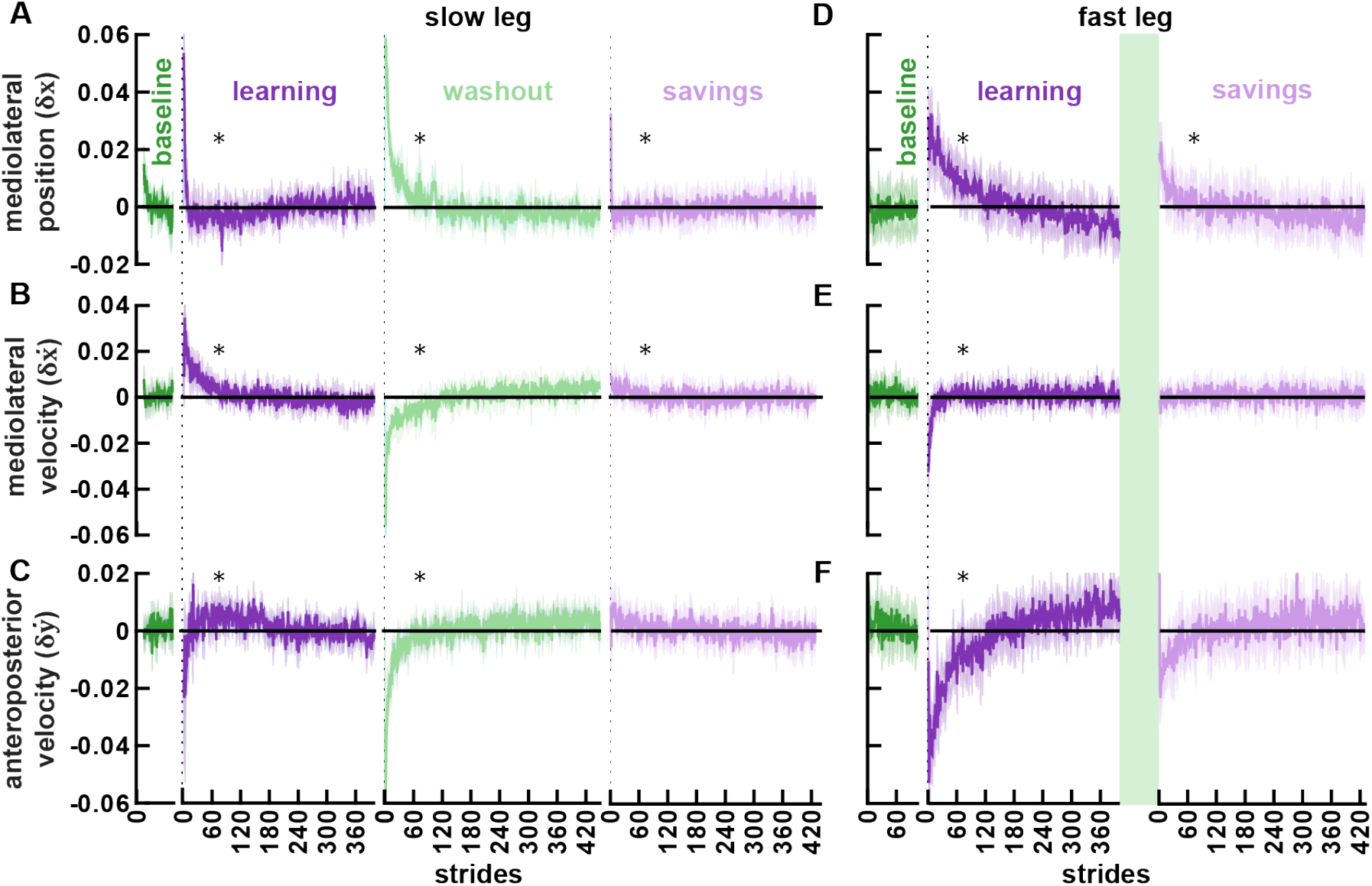
Centre of mass error trajectories. Mean and standard error are depicted with solid lines and shading, respectively. Black asterisks indicate trajectories for which the early error is significantly different than late error. For the slow leg (A-C), error magnitude in early learning and early washout was significantly higher than later in the respective blocks, for all states. Savings error was significantly higher for δx. In the fast leg (D-F), early CoM error was significantly higher for each state during the learning block, and for δx during the savings block. Data presented here are demeaned for clarity of presentation; statistical tests (Section 2.3) were conducted on raw data.

The same trends appear in the washout block except for the initial 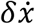, which shows strong aftereffects. The sign of the error is reversed from the initial error in the learning block, with a positive, laterally-directed velocity error in the fast leg and a negative, medially-directed error in the slow leg. The savings block shows similar trends as the learning block, but with lower initial errors in all states, for both legs. Overall, CoM state error was reduced across phases, pointing towards its relevance as a measure of performance and providing a foundation for further analyses.

### 3.2 A linear model captures meaningful stride-to-stride CoM error dynamics

Having established that CoM error is reduced during learning and washout, we confirmed stability of the linear dynamics and consistent goodness of fit for our model-based approach (NRMSE < 10%, Supplement S6). We validate the model’s relative predictive capabilities using K-step skill scores relative to benchmark models.

Per block and for both legs, the average skill score across states is positive for at least three strides, indicating that the model outperforms both the persistence and mean predictor benchmarks (Figure 4). Nontrivial error dynamics are thus consistently captured by the model over a short time horizon. The mean-skill approaches zero in all blocks, as expected for a stable system, while the persistence-skill becomes negative as the persistence model eventually outperforms the error model in baseline and washout.

**Figure 4:**
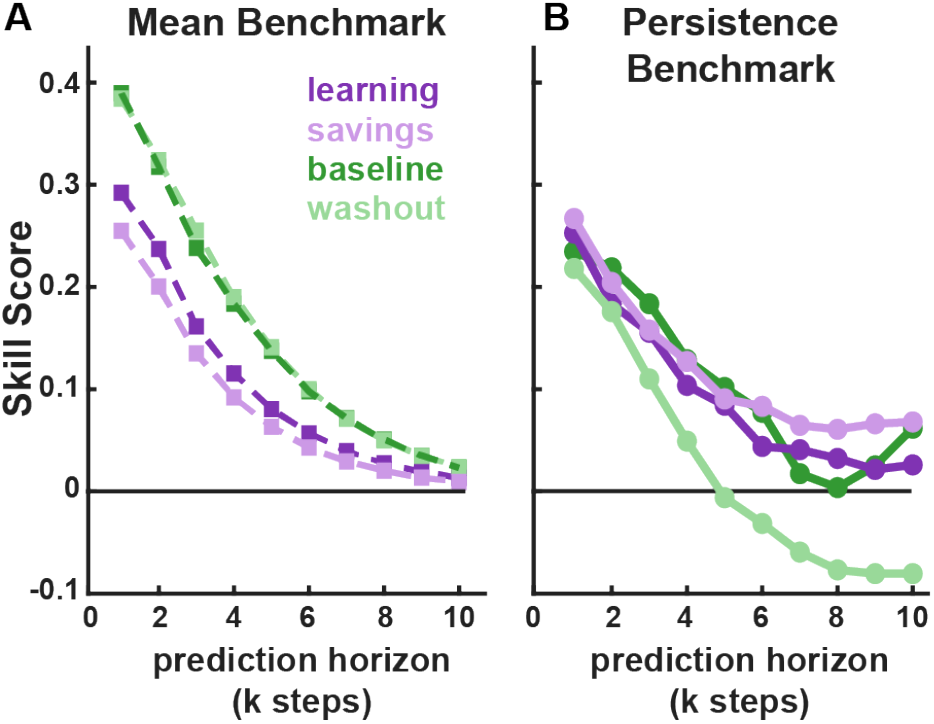
Per-block slow leg average skill across all three error states, with respect to the mean-predictor benchmark (A) and the persistence benchmark (B). A positive score indicates that the CoM error map outperforms the benchmark model at the k^th^ step.

For the slow leg, the average single step persistence skill score per block ranged from 0.22 to 0.27, while the average skill score relative to the mean predictor ranged from 0.25 to 0.39 and is comparable to *R*^2^ values reported in similar prior work (Supplement S6). We interpret the persistence scores being consistently above 10% as sufficient improvement over this benchmark due to the high degree of autocorrelation inherent to gait data (per-state, per-leg skill scores in Supplement S6).

### 3.3 Steady-state error dynamics differ significantly between experimental blocks

After confirming CoM error as a relevant learning signal and validating our stride-to-stride error model, we compared the error dynamics between the baseline, learning, washout, and savings blocks.

#### 3.3.1 Stability

To compare the stability of different blocks, we computed a bootstrapped distribution of the largest eigenvalue for each block and for each leg. The 95% confidence interval of the real part of the eigenvalue (spectral radius *ρ*(*A*_block_)) indicates that the linear dynamics are stable and the largest eigenvalue is real across all blocks, for both legs (Table 1). In all blocks, the confidence intervals are shifted lower for the slow leg than for the fast leg. This increased stability may be simply attributable to the reduced belt speed. Within each leg, the intervals for each block strongly overlap. This indicates little change in steady-state stability due to the split-belt perturbation, at least as captured by this scalar metric.

**Table 1:** Largest eigenvalue by block and leg (95% Confidence Interval)

| Block | Slow Leg | Fast Leg |
| --- | --- | --- |
| B (slow) | 0.72 (0.63–0.83, $\pm 0.02i$) | —* |
| B (fast) | —* | 0.87 (0.80–0.92, $\pm 0.00i$) |
| L | 0.70 (0.60–0.76, $\pm 0.00i$) | 0.88 (0.82–0.92, $\pm 0.00i$) |
| W | 0.74 (0.67–0.81, $\pm 0.02i$) | —* |
| S | 0.67 (0.55–0.76, $\pm 0.00i$) | 0.85 (0.75–0.91, $\pm 0.00i$) |

\* Slow baseline data analysed only for leg assigned to slow belt during split-belt, no data analysed for fast-leg washout.

#### 3.3.2 Eigenstructure

We then used a contrast analysis to quantify the similarity between eigenvector direction across blocks, finding observable changes in their orientation (Figure 5). For the slow leg, the contrast analysis (Equation 2.4) indicated that the dominant directions of the savings and learning dynamics are significantly more similar to each other than either is to the baseline dominant subspace (d_slow_ = 0.39,95% CI = [0.02,0.80]). The same contrast was not significant for the fast leg (d_fast_ = 0.05,95% CI = [−0.04,0.17]) . Thus while the eigenvalues during learning and savings may be similar to baseline, the dynamics in learning and savings are fundamentally distinct from the dynamics in baseline.

**Figure 5:**
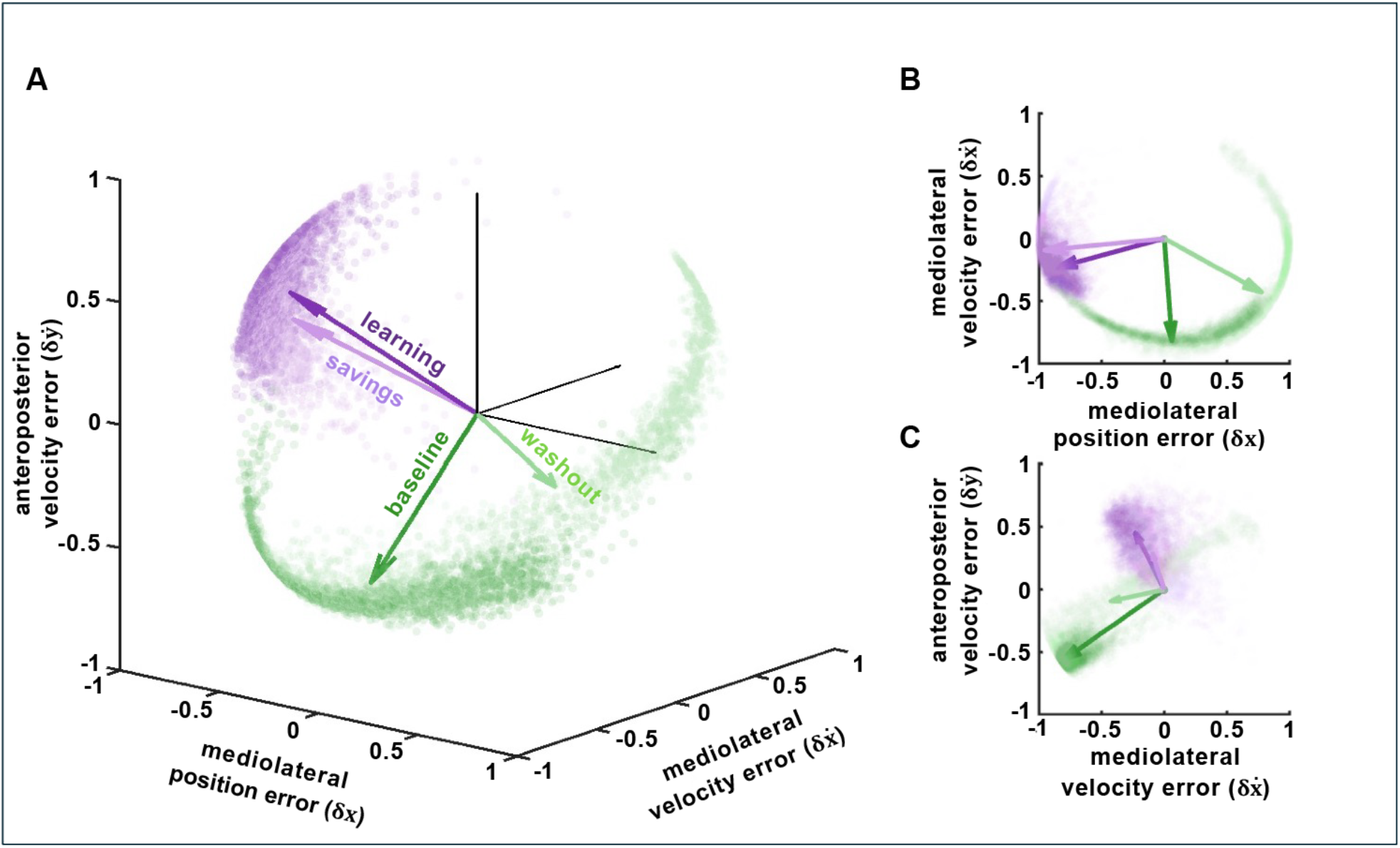
Eigenvectors from the CoM error map for baseline (dark green), washout (light green), learning (dark purple), and savings (light purple) alongside the bootstrapped distribution (purple and green dots). Bootstrapped data from the split-belt blocks cluster together, as do the noisier tied-belt data (A). Rotation away from the 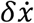 direction in the split-belt blocks relative to the tied-belt blocks is particularly visible in the 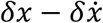 frontal plane (B) and the 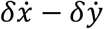 velocity plane (C).

Next we examined the similarity between belt conditions, and asked whether the tied belt conditions were more similar to each other than split-belt conditions, and vice versa. The tied-belt vs split-belt contrast (Equation 2.5) for the slow leg was also significant, indicating that dynamics arising from the same belt condition are more similar to each other than to those from the opposing condition (d_cluster_ = 0.32,95% CI = [0.05,0.61]).

The change in orientation of the dominant eigenvectors for the different slow-leg experimental blocks is visualized in Figure 5. The direction of longest error persistence is consistently shifted away from mediolateral velocity during split-belt conditions. In the velocity plane (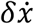 vs 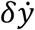), the dominant learning eigenvector is rotated away from the mediolateral velocity direction 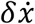 and more closely aligned with 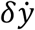. In the mediolateral plane (*δx* vs 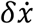), the vector is rotated toward the mediolateral *δx* position axis and away from 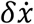. Thus, during tied-belt walking, mediolateral velocity error dominates, while during split-belt walking, error persists in directions that are almost orthogonal to mediolateral velocity error.

#### 3.3.3 Post-hoc analysis of fastest eigenvectors

The shift in the direction of the dominant eigenvector indicates a possible change in control priorities away from anteroposterior velocity error and toward mediolateral velocity error. To explore whether participants in fact reduce mediolateral velocity error faster during split-belt walking, we conducted a post-hoc analysis of the fastest eigenvectors between blocks. The fastest eigenvector is defined as the vector corresponding to the smallest eigenvalue. We analysed our results using the same statistical and bootstrapping approach that we used to analyse the dominant eigenvectors in Section 2.5. *A*_block_ matrices computed from raw data show that the fastest eigenvector is indeed rotated *towards* the mediolateral velocity axis, further supporting this hypothesis, and bootstrapped results found that the fastest eigenvector of learning and savings is significantly different from baseline 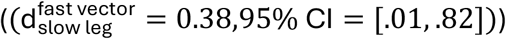. On the other hand, d_cluster_ was not significantly different for the fast eigenvector comparison.

### 3.4 Learned structure explains early savings CoM error trajectories

To explore whether transient error reduction during savings reflects the dynamical structure of the learning block, we compared how well the steady state dynamics from learning and washout could predict the first minute of the savings block. We used the first 30 strides of the savings block and computed the skill score of the steady state learning model (*A*_L_) relative to the washout model (*A*_W_) over 2000 bootstrap iterations, resampling by participant to mitigate the effect of outliers. The learning model exhibits higher skill in predicting frontal and sagittal plane velocity error, but not in mediolateral position, where the bootstrapped results do not favour one model over the other 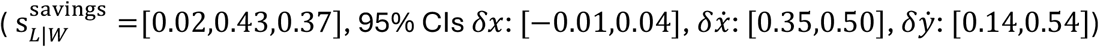.

We also considered that the superior ability of the learning dynamics relative to washout dynamics to explain savings may be because the error maps reflect the constraints of split-belt and tied-belt walking. To determine whether our results were reflecting the mechanical constraints rather than adaptation, we compared the ability of the steady-state learning (*A*_L_) to predict early learning (first 30 strides of the learning block) with its predictive power for the first 30 strides of early savings. Here, we find that the dynamical structure of steady-state learning (*A*_L_) explains significantly more of early savings than early learning, in every state 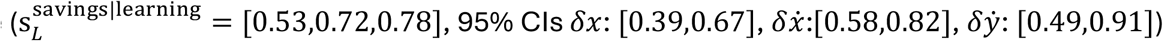.

Taken together, these results show that early savings reflects the dynamical structure of the learning block. Importantly, as error correction during early savings is better predicted by *A*_L_ than error correction during early learning, the similarity to the steady state learning dynamics is not just an artifact of split-belt constraints. Since these dynamics are not immediately imposed during early learning, when participants are first exposed to the split-belt, this finding suggests that learned dynamics are rapidly recalled during savings.

## 4 Discussion

Our results show that as people adapt, they reduce CoM error in the mediolateral and anteroposterior directions, and that locomotor adaptation can significantly alter the structure of CoM error dynamics, persistently reorganizing the geometry of CoM movement regulation. This reorganization is not immediately apparent from the scalar results, but rather is revealed through the significant similarity of the linear error dynamics in the respective experimental conditions, i.e. tied-belt vs split-belt. The steady state dynamics learned during the first exposure to the treadmill can predict the early transient response upon re-exposure to the split-belt, providing further evidence that a new dynamical structure is not only learned, but may also be recalled.

These major changes in the structure of the CoM error dynamics are detected by analysing the direction of the eigenvector corresponding to the dominant eigenvalue for each experimental block (baseline, learning, washout, savings). Scalar metrics such as the margin of stability are not sufficient to clarify these significant changes, nor are spatiotemporal outcomes such as step length asymmetry. The spectral radius also did not reveal between-block differences, although similar stability metrics are used quite frequently to quantify gait stability [30]. Assessing modifications in stability regulation from the geometric perspective of eigenvector direction is not just more sensitive to differences between blocks, but also provides a mechanistic explanation as to how the CoM error dynamics shift in space, indicating which error directions are prioritized or de-prioritized after locomotor adaptation.

The insights gleaned from the eigenvector analysis also point to adaptation being coordinated via the dynamics of the slow leg. While both legs exhibit scalar error reduction, the observed changes in steady-state error dynamics between the tied belt (baseline and washout) blocks and the split-belt (learning and savings) blocks are concentrated in the slow leg, with no significant between-block differences found in the fast leg. These findings are consistent with prior work on spatiotemporal adaptation measures that show changes in slow-leg kinetics and kinematics drive changes in asymmetry measures [1], [27], [31]. The changes in eigenstructure suggest stability is regulated asymmetrically, and primarily by the slow leg. However, more targeted experiments are needed to confirm this, such as studies where participants experience systematic CoM perturbations at slow and fast midstance.

Geometrically, velocity in the sagittal plane is associated with the direction of slowest error decay during split-belt walking. During tied-belt walking, error decays the slowest in the direction of mediolateral velocity, particularly during baseline. Prior work has shown that forward velocity is actively regulated during nominal tied belt walking [21], corresponding to the constraint of not walking off the treadmill. Taken together, this implies that during split-belt walking, people de-prioritize CoM error in in this direction, possibly prioritizing mediolateral velocity error. The results of our post hoc analysis of the fastest eigenvalues provide evidence of prioritization of mediolateral velocity error, showing that the fastest eigenvector is more aligned in this direction during split-belt conditions than during baseline. Significant geometric changes in both the dominant (slow) eigenstructure and in the fast eigenstructure point to a shift in control priorities, altering how different components of CoM motion dominate and contribute to stride-to-stride stabilization during adaptation.

The structural changes observed during steady-state split-belt walking are also reflected in early strides of the savings block. Although the steady-state dynamics of learning and savings are not identical, the learning dynamics predict the early strides of savings and specifically predict velocity in the frontal and sagittal planes significantly better than the more recently-experienced washout dynamics. The predicted mediolateral error trajectory for early savings is nearly identical between the washout and learning models (Figure 6A), as *A*_L_ and *A*_W_ encode similar mediolateral position dynamics (matrix entries reported in Supplement S8). In contrast, the velocity dynamics in both the frontal and sagittal plane differ more substantially between the two models, resulting in clearer differences in their predictions (Figures 6B and 6C).

**Figure 6:**
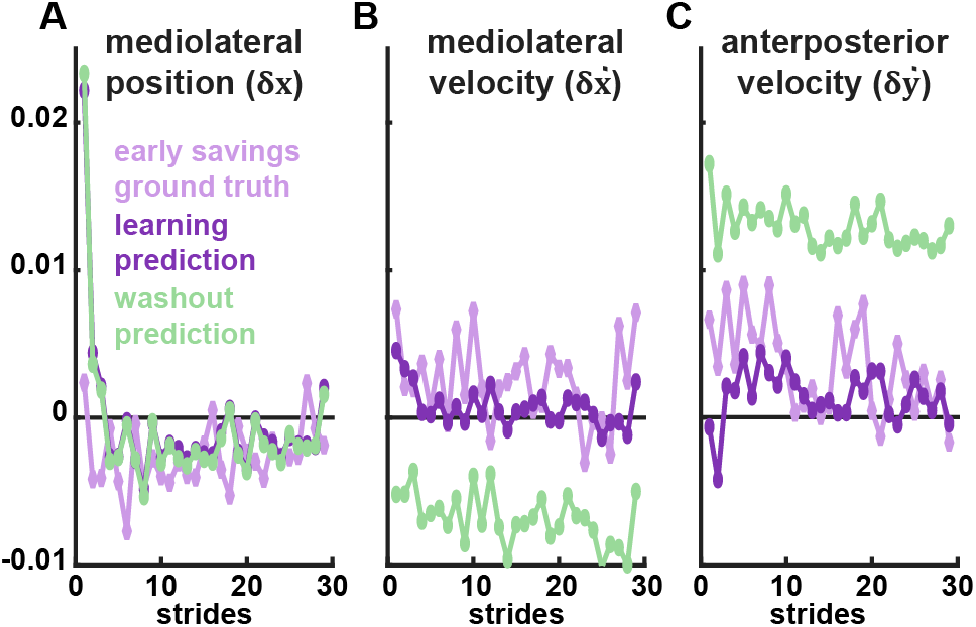
CoM error during the first thirty strides of the slow leg in the savings block (ground truth, pale purple with diamond markers), alongside the learning model (A_L_, dark purple) and washout model (A_W_, light green) predictions.

The observed similarity between learning and savings, both in the eigenstructure of the steady-state matrices and in the dynamics of early savings, could merely reflect the novel mechanical constraints of split-belt walking. However, our results show that the steady-state learning dynamics predict early savings significantly better than early learning. Taken together, this suggests that the altered CoM error regulation observed between tied- and split-belt conditions is not instantaneously imposed by the split-belt but is actively learned. While preliminary, our findings suggest that the structure of this altered regulation strategy may be rapidly recalled upon re-exposure to the split-belt during savings.

Based on the low initial error in the slow leg savings block (Figure 3D-F), there seems to be an almost instantaneous re-engagement of the learned split-belt dynamics, but more work is needed to study confirm existence of this effect. It is possible that savings is best explained as a combination of washout and learning dynamics. Individual differences are also likely to be present, with some individuals defaulting to their baseline strategy, some remaining in the washout regime, and some individuals exhibiting stronger recall.

Recent work has proposed that locomotor learning may include mechanisms of recall that enable rapid re-adaptation [12], [32]. The approach shown here provides a useful parameterization for continuing to probe this question. With respect to non-scalar metrics, our approach aligns with recent work showing that postural control is reorganized geometrically in new contexts [33]. Our results contribute to a growing body of evidence showing that stability-relevant adaptations are present during locomotor adaptation and learning [1], [5], [34], [35], [36], [37].

Across participants, scalar CoM error was consistently higher during early learning and early washout, particularly mediolateral CoM error. This is even though the perturbation occurred in the sagittal plane, aligning with findings from [25] showing that CoM error regulation is coupled across dimensions during perturbed nominal walking. The sudden asymmetry induced in the gait dynamics may further contribute to this effect.

The linear error dynamics models exhibit reasonable performance and consistently low NRMSE. With respect to the skill score, as noted, we consider the persistence baseline to be more informative for our model. This score is fairly consistent across blocks and states, decaying with increasing number of strides as expected. Although there is no direct comparison in the literature, our model performance with respect to the mean-predictor benchmark, i.e. *R*^2^ is comparable to prior work (Supplement S6).

The method presented here is experimentally feasible, using only a minimal marker set and not requiring explicit separation of the plant and controller. Our high-level view of dynamics at the stride-to-stride level is also a limitation. While we have shown interesting changes in the overarching closed loop dynamics of stride-to-stride error regulation, our current analysis does not explicitly model control. Thus, we can only describe the downstream effects of locomotor adaptation, not the explicit changes in control strategy driving the observed changes. We are also limited by the use of pooled data, and our results should be interpreted as systems-level effects. As with any human behaviour, there are likely crucial individual differences in CoM error regulation and adaptation, but constructing these models requires hundreds of strides. Explicitly modelling changes in error dynamics on an individual basis and at a shorter timescale would reveal the actual process by which these dynamics are adapted. We plan to continue this work along these lines, seeking ways to develop individualized models of adaptation and applying these models to shorter trajectories.

## 5 Conclusions

We proposed a novel, dynamical systems approach for analysing stability-relevant locomotor adaptations to novel walking environments. Using this framework, we show that walking on a split-belt treadmill alters the geometric structure of CoM movement regulation, substantially changing the direction in which CoM error decays the slowest and dissipates the fastest. These changes persist after a washout period and are recalled rapidly upon re-exposure during savings. Scalar approaches to quantifying gait stability, even those based on dynamical systems theory, cannot detect these changes. The precise evolution of CoM error dynamics from initial exposure through savings remains an open question and merits further study.

## Acknowledgment

We thank Varun Joshi for his helpful comments.

## Notes

### Competing Interest Statement

The authors have declared no competing interest.

### Funding

- U.S. National Science Foundation: CAREER award 1352632, DGE 2040434
- National Institutes of Health: 1R01NS096083

