## Supplement for "Locomotor adaptation can persistently reorganize stride-to-stride regulation of centre-of-mass error dynamics"

July 2026

#### S1 Experiment

We use a previously collected dataset consisting of motion tracking data for 21 participants learning to walk on a split-belt treadmill with different belt speeds [1]. Pelvic (anterior and posterior superior iliac spines) and ankle marker (malleolus) data were available for 15 of these 21. One participant was excluded due to their raw marker data containing substantial tracking artifacts. The final analysis therefore included 14 participants (9 female, age $24 \pm 3.6$ years, weight $71 \pm 15.2$ kg, height $1.64 \pm 0.13$ m). Raw marker data is available **here** .

#### S2 Data processing

Marker data are recorded in a stationary global frame fixed to the treadmill and is processed as follows. We approximate the centre of mass (CoM) as the average of the four pelvic markers (Left/Right Posterior/Anterior superior iliac spines), numerically computing the derivative of this signal to get the CoM velocity. We sample CoM data at midstance, which we define as the moment when the CoM is aligned with the stance foot in the sagittal plane. This is approximated with marker data as the time points in the gait data where the averaged CoM position is aligned with the ankle marker and has forward velocity. The ankle marker position and velocity define the reference state, and we define CoM error with respect to this reference. Thus, given set $K$ of time indices corresponding to midstance, we compute the CoM error state relative to the foot:

$$\delta\mathbf{x}[k] = \begin{bmatrix} x_{\text{ankle}}[k] - x_{\text{CoM}}[k] \\ \dot{x}_{\text{ankle}}[k] - \dot{x}_{\text{CoM}}[k] \\ \dot{y}_{\text{ankle}}[k] - \dot{y}_{\text{CoM}}[k] \end{bmatrix} \tag{1}$$

where $k \in K$. All position variables are non-dimensionalized per participant by dividing by leg length $l$ and velocity variables by $\sqrt{g \cdot l}$. The error trajectories are processed and computed separately for each of the experimental blocks (Baseline, Learning, Washout, Savings) and for the right and left legs. An example script demonstrating our methodology is available **here** and will be uploaded to a public repository upon publication.

### S3 Fitting CoM error dynamics models

Once the midstance CoM error states at each stride are computed, we can fit $A_{\text{block}}$ for each experimental block. As we wish to model steady-state behaviour, we discard the first thirty seconds of the slow and fast baseline error trajectories, and we discard the first minute of the learning, washout, and savings trajectories. Due to the differing durations of the experimental blocks, this leaves one and a half minutes of data for the baseline block, nine minutes for the learning and savings blocks, and fourteen minutes for the washout block. We select one minute based on prior work showing that the margin of stability, which is a linear combination of CoM position and velocity, converges in under one minute [2]. We additionally confirmed that for two, three, and four minute-length trims of the initial portion of the non-baseline blocks, our conclusions remain the same.

We pool the trimmed data across participants and demean the resulting trajectories so that the estimated steady-state operating point is centered at the origin. We transform the mediolateral position and velocity errors such that position error lateral to the CoM is positive. For each block, we seek to compute the matrix $A_{\text{block}}$

$$\delta \mathbf{x}(n+1) = A_{\text{block}} \delta \mathbf{x}(n) \tag{2}$$

where block $\in$ {B,L,W,S}. Solving for $A_{\text{block}}$ amounts to a least squares problem:

$$A_{\text{block}} = \arg\min_{A} \sum_{n=1}^{N-1} \left\| \delta \mathbf{x}_{\text{block}}(n+1) - A \, \delta \mathbf{x}_{\text{block}}(n) \right\|_2^2 , \tag{3}$$

For each experimental block, we construct concatenated data matrices $X_{\text{block}}$ and $Y_{\text{block}}$ whose rows contain CoM error states at midstance $\delta \mathbf{x}_{\text{block}}(n)^T$ and their corresponding successor states at the next midstance $\delta \mathbf{x}_{\text{block}}(n+1)^T$, respectively. We estimate the dynamics matrix $A_{\text{block}}$ by solving

$$A_{\text{block}}^T = \arg\min_{\Theta} \left\| Y_{\text{block}} - X_{\text{block}} \Theta \right\|_F^2 . \tag{4}$$

The ordinary least-squares solution is

$$A_{\text{block}}^T = X_{\text{block}}^{\dagger} Y_{\text{block}}, \tag{5}$$

where subscript $F$ denotes the Frobenius norm and superscript $\dagger$ denotes the Moore-Penrose pseudo-inverse.

#### S4 Model validation

We confirm that the resulting model is plausible, has reasonable predictive power, and captures a genuine signal of interest, using three complementary approaches to validate our linear error dynamics. We first check the physical plausibility of the discrete-time model by computing the spectral radius $\rho(A_{\text{block})} = \max|\lambda_i| < 1$ for all blocks, where $\lambda_i$ is the $i$th eigenvalue. We confirm that $\rho(A_{\text{block})} < 1$, ensuring that each linear map is stable. We then assess predictive accuracy by computing the root mean squared error (RMSE), normalized by the range of values in the data. We compute this normalized root mean squared error (NRMSE) per leg for each CoM error state, per block. We compute the one step prediction error only, and expect that this error should be less than 20% for a sufficiently useful model. Skill scores are computed from MSE aggregated across all valid stride pairs $(\delta x(n), \delta x(n+k))$, where $n = 1, \ldots, N-k$. For $k > 1$, model predictions are generated by iterating the map k times:

$$\delta\hat{\mathbf{x}}(n+k) = A^k\,\delta\mathbf{x}(n) \tag{6}$$

The persistence benchmark model predicts the current state vector over horizon $k$

$$\delta\hat{\mathbf{x}}(n+k) = \delta\mathbf{x}(n), \tag{7}$$

while the mean predictor benchmark model, essentially a zero-predictor, predicts the sample mean:

$$\delta\hat{\mathbf{x}}(n+k) = \delta\bar{\mathbf{x}}. \tag{8}$$

#### S5 Comparing model dynamics

Our analysis of dynamic similarity between blocks is based on comparing the directions of the dominant eigenvectors of the different $A_{\text{block}}$ matrices, repeating the analysis over 2000 bootstrap iterations with participant-level resampling. If the dominant eigenvalue is complex, the two largest eigenvalues are a complex conjugate pair and we instead use the two-dimensional subspace spanned by the real and imaginary components of the corresponding vectors. Likewise, if the ratio of the magnitudes of the second-largest and largest eigenvalues exceeds 0.85, indicating that there is no clearly dominant one-dimensional eigenspace, we compare the dominant two-dimensional subspace spanned by the two leading eigenvectors. Similarity is then quantified either by the maximum principal angle between subspaces, measuring the worst-case directional misalignment, or by computing the magnitude of the projection of one block's dominant eigenvector onto the dominant subspace of the comparison block. Like the dominant eigenvector, the dominant subspace represents the directions of slowest error decay, along which system error persists the longest. Mean pairwise similarity indices and 95% confidence intervals for all within- and between-block comparisons are shown in Figure 1, alongside the corresponding bootstrapped contrast statistics (Equations 2.4 and 2.5).

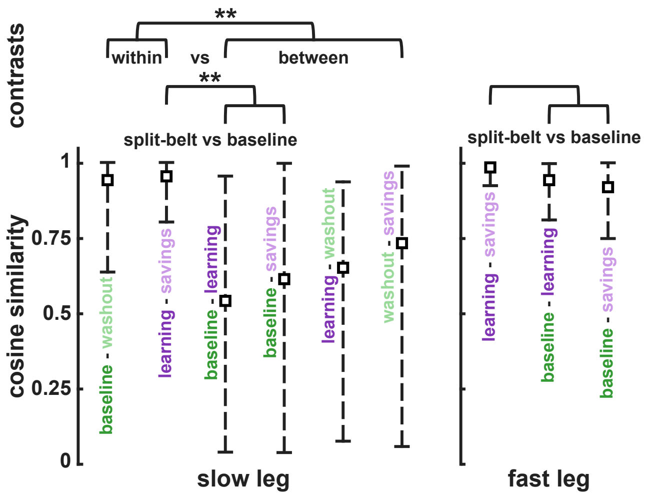


Figure 1: Eigenstructure Similarity Analysis. Bootstrapped means and 95% confidence intervals of the pairwise comparisons used in the contrast analyses (Equations 2.4 and 2.5 in the main text). Significant contrasts are marked by a double asterisk at the top of the figure.

### S6 Validation results

#### S6.1 Spectral radius and NRMSE

For all tables in this section, note that data from the slow baseline and washout blocks were not analysed for the assigned fast leg, and fast baseline data were not analysed for the assigned slow leg. This is indicated on the tables with a '—'.

The estimated maps were stable across all blocks and legs, with spectral radii $\rho(A_{\mathrm{blk}}) < 0.74$ for the slow leg and $\rho(A_{\mathrm{blk}}) < 0.88$ for the fast leg (Table 1), indicating biological plausibility of the learned error dynamics.

Table 1: Spectral radius $\rho(A_{\mathrm{block}})$ by block and leg.

| **Block** | **Slow Leg** | **Fast Leg** |
|---|---|---|
| Baseline (slow) | 0.69 | — |
| Baseline (fast) | — | 0.88 |
| Learning | 0.70 | 0.88 |
| Washout | 0.74 | — |
| Savings | 0.67 | 0.85 |

Normalized RMSE was consistent across all states and blocks, ranging narrowly from 0.04 to 0.12 and well below our 20% benchmark (Table 2).

Table 2: NRMSE by block and state variable for each leg.

| **Leg** | **State** | **Block NRMSE** | | | | |
|---|---|---|---|---|---|---|
| | | ***B (slow)*** | ***B (fast)*** | ***L*** | ***W*** | ***S*** |
| Fast | $x$ | — | 0.08 | 0.08 | — | 0.09 |
| | $\dot{x}$ | — | 0.09 | 0.10 | — | 0.09 |
| | $\dot{y}$ | — | 0.05 | 0.05 | — | 0.04 |
| Slow | $x$ | 0.09 | — | 0.09 | 0.06 | 0.09 |
| | $\dot{x}$ | 0.12 | — | 0.10 | 0.07 | 0.10 |
| | $\dot{y}$ | 0.09 | — | 0.07 | 0.07 | 0.06 |

#### S6.2 Skill scores

Skill scores are reported below. The persistence predictor is our primary benchmark, as positive skill indicates predictive information beyond stride-to-stride autocorrelation. We interpret persistence skill scores greater than 0.10 as evidence that the model captures meaningful stride-to-stride dynamics, and indeed this outcome is fairly across blocks and states.

Table 3: Single-step skill scores for each state variable.

**(a) ML position**

| | **Slow Leg** | | **Fast Leg** | |
|---|---|---|---|---|
| **Block** | **Pers.** | **Mean** | **Pers.** | **Mean** |
| Baseline (slow) | 0.17 | 0.43 | – | – |
| Baseline (fast) | – | – | 0.08 | 0.79 |
| Learning | 0.17 | 0.47 | 0.12 | 0.72 |
| Washout | 0.16 | 0.48 | – | – |
| Savings | 0.16 | 0.46 | 0.11 | 0.70 |

**(b) ML velocity**

| | **Slow Leg** | | **Fast Leg** | |
|---|---|---|---|---|
| **Block** | **Pers.** | **Mean** | **Pers.** | **Mean** |
| Baseline (slow) | 0.20 | 0.48 | – | – |
| Baseline (fast) | – | – | 0.24 | 0.32 |
| Learning | 0.31 | 0.16 | 0.23 | 0.37 |
| Washout | 0.17 | 0.47 | – | – |
| Savings | 0.33 | 0.12 | 0.11 | 0.46 |

**(c) AP velocity**

| Block | Slow Leg Pers. | Slow Leg Mean | Fast Leg Pers. | Fast Leg Mean |
|---|---|---|---|---|
| Baseline (slow) | 0.33 | 0.26 | – | – |
| Baseline (fast) | – | – | 0.19 | 0.30 |
| Learning | 0.28 | 0.25 | 0.25 | 0.47 |
| Washout | 0.32 | 0.20 | – | – |
| Savings | 0.31 | 0.18 | 0.11 | 0.49 |

We are not aware of any split-belt studies to which we can reasonably compare our model performance. Our model is meant to predict local fluctuations in the error dynamics, and we cannot expect similar performance to studies using high-magnitude perturbations to model reactive balance recovery such as [3]. However, although it is not a direct comparison to stride-to-stride analysis, $R^2$ values for step-to-step maps computed for CoM error fluctuations during nominal tied-belt walking have been reported [4]. Comparing our single-step mean-skill scores ($R^2$) for the slow and fast leg during the tied-belt baseline and washout blocks, our scores are comparable to these step-to-step $R^2$ values in [4]. The sole exception is the single-step mean-skill score for $\delta x$ in the slow leg (our skill score $\approx 0.45$, their $R^2 \approx 0.80$).

This reduced performance with respect to $R^2$ possibly because the CoM position error relationship is not as strong at the stride-to-stride level, or because our choice of slow speed is much lower than the speeds used for nominal tied belt walking in [4] (0.5 m/s vs an average of 1.2 m/s). Across all states, the one-step mean-skill score for the fast leg is more comparable to prior work, particularly in the baseline tied-belt block. We further note that single-step $R^2$ values are lower for mediolateral velocity error $\delta\dot{x}$ in the split-belt blocks than in tied-belt blocks, however the persistence skill score is higher for the split-belt blocks.

### S7 RM-ANOVA and Planned comparisons

RM-ANOVA and planned comparisons between Early/Late Baseline (EB/LB), Early/Late Learning (EL/LL), Early/Late Washout (EW/LW), and Early/Late Savings (ES/LS).

Table 4: Repeated Measures ANOVA Results

| Outcome | Leg | *df* | *F* | *p* |
|---|---|---|---|---|
| ML Pos | Slow | (7, 91) | 24.30 | $2.42 \times 10^{-18}$ |
| | Fast | (5, 65) | 18.93 | $1.43 \times 10^{-11}$ |
| ML Vel | Slow | (7, 91) | 23.22 | $1.009 \times 10^{-17}$ |
| | Fast | (5, 65) | 25.74 | $3.16 \times 10^{-14}$ |
| AP Vel | Slow | (7, 91) | 10.11 | $2.68 \times 10^{-9}$ |
| | Fast | (5, 65) | 5.83 | $1.66 \times 10^{-4}$ |

Table 5: Planned pairwise $t$-test results for each state variable.

**(a) ML position**

| **Comparison** | **Leg** | $\boldsymbol{t}$ | $\boldsymbol{p}$ |
|---|---|---|---|
| EL vs LL | Slow | 5.12 | 0.004 |
| | Fast | 5.65 | $1.60 \times 10^{-4}$ |
| EW vs LW | Slow | 8.22 | $4.94 \times 10^{-6}$ |
| | Fast | —* | —* |
| ES vs LS | Slow | 2.21 | 0.046 |
| | Fast | 4.98 | $2.53 \times 10^{-4}$ |

**(b) ML velocity**

| **Comparison** | **Leg** | $\boldsymbol{t}$ | $\boldsymbol{p}$ |
|---|---|---|---|
| EL vs LL | Slow | 5.31 | $2.90 \times 10^{-4}$ |
| | Fast | −5.52 | $1.97 \times 10^{-4}$ |
| EW vs LW | Slow | −5.69 | $2.20 \times 10^{-4}$ |
| | Fast | —* | —* |
| ES vs LS | Slow | 3.03 | 0.01 |
| | Fast | −1.93 | 0.08 |

**(c) AP velocity**

| **Comparison** | **Leg** | $\boldsymbol{t}$ | $\boldsymbol{p}$ |
|---|---|---|---|
| EL vs LL | Slow | −1.16 | 0.343 |
| | Fast | −4.14 | 0.0023 |
| EW vs LW | Slow | −6.84 | $3.59 \times 10^{-5}$ |
| | Fast | —* | —* |
| ES vs LS | Slow | 1.45 | 0.343 |
| | Fast | −0.55 | 0.616 |

* No data were analysed for fast-leg washout.

### S8 Linear models ($A_{\mathbf{block}}$ matrices)

Slow leg linear maps:

$$A_{\mathrm{L}}^{\mathrm{slow}} = \begin{bmatrix} 0.6669 & 0.0803 & -0.0066 \\ 0.0736 & 0.3661 & -0.0264 \\ -0.1110 & -0.0729 & 0.4425 \end{bmatrix}$$

$$A_{\mathrm{S}}^{\mathrm{slow}} = \begin{bmatrix} 0.6771 & 0.0644 & 0.0097 \\ 0.0154 & 0.3393 & -0.0616 \\ -0.0769 & -0.0818 & 0.3941 \end{bmatrix}$$

$$A_{\mathrm{W}}^{\mathrm{slow}} = \begin{bmatrix} 0.7098 & 0.0093 & 0.0413 \\ -0.0379 & 0.6217 & 0.1109 \\ 0.0622 & 0.2049 & 0.3168 \end{bmatrix}$$

$$A_{\mathrm{B}}^{\mathrm{slow}} = \begin{bmatrix} 0.6672 & 0.0210 & -0.0340 \\ -0.0488 & 0.5837 & 0.1683 \\ 0.0105 & 0.2779 & 0.3083 \end{bmatrix}$$

Fast leg linear maps:

$$A_{\mathrm{L}}^{\mathrm{fast}} = \begin{bmatrix} 0.8198 & 0.1407 & -0.0962 \\ -0.0144 & 0.5136 & 0.1075 \\ -0.1846 & 0.2416 & 0.5388 \end{bmatrix}$$

$$A_{\mathrm{S}}^{\mathrm{fast}} = \begin{bmatrix} 0.8050 & 0.1131 & -0.0951 \\ -0.0169 & 0.5730 & 0.1153 \\ -0.1797 & 0.2644 & 0.5289 \end{bmatrix}$$

$$A_{\mathrm{B}}^{\mathrm{fast}} = \begin{bmatrix} 0.8679 & 0.0766 & -0.0758 \\ -0.0105 & 0.5115 & 0.0676 \\ -0.0829 & 0.0238 & 0.7254 \end{bmatrix}$$
